# Quantitative analysis of Histone modifications in gene silencing

**DOI:** 10.1101/2020.11.17.386896

**Authors:** Kenneth Wu, Namrita Dhillon, Kelvin Du, Rohinton T. Kamakaka

## Abstract

Gene silencing in budding yeast is mediated by Sir protein binding to unacetylated nucleosomes to form a chromatin structure that inhibits transcription. This transcriptional silencing is characterized by the high-fidelity transmission of the silent state. Despite its relative stability, the constituent parts of the silent state are in constant flux giving rise to a model that silent loci can tolerate such fluctuations without functional consequences. However, the level of tolerance is unknown and we developed a method to measure the threshold of histone acetylation that causes the silent chromatin state to switch to the active state. We show that loss of silencing required between 50% and 75% of the unacetylated histones to be replaced with acetylated histone mimics. The precise levels of unacetylated nucleosomes required varied from locus to locus and was influenced by both silencer strength and UAS enhancer/promoter strength. Simple calculations suggest that an approximately 50% reduction in the ability of acetylases to acetylate individual nucleosomes across a large domain may be sufficient to generate a transcriptionally silent region in the nucleus.

## Introduction

Multiple loci in yeast are transcriptionally silenced including the cryptic mating type loci *HML* and *HMR* on chromosome III as well as sub-telomeric sites (Gartenberg and Smith, 2016). At *HML* and *HMR*, DNA elements called silencers serve as binding sites for specific proteins, which in turn recruit the repressor proteins Sir1, Sir2, Sir3 and Sir4 (Chien et al., 1993; Fox et al., 1997; Hecht et al., 1996; Liu and Lustig, 1996). The histones at silent loci lack acetylation or methylation marks (O’Kane and Hyland, 2019) though they are enriched in phosphorylated histone H2A (Kirkland and Kamakaka, 2013; Kitada et al., 2011). The Sir2/Sir4 heterodimer deacetylates K9 in histone H3 and K16 in histone H4 thereby facilitating Sir3 binding to the unmodified histone tails (Luo et al., 2002; Rusche et al., 2002). Sir3, in turn, simultaneously interacts with and stabilizes the binding of the Sir2/Sir4 heterodimer with nucleosomes thus generating a feedback loop that aids in further binding and spreading of the Sir proteins along the nucleosomal filament (Gartenberg and Smith, 2016). Sir protein interactions with nucleosomes hinder the association of the transcription machinery with regulatory sequences thereby establishing the transcriptionally silent state at *HML* and *HMR.*

Besides the silencers and the Sir proteins, the post-translational modifications of the histones play a critical role in silencing. Studies utilizing various histone mutants have shown that a region of the histone H4 N-terminal tail from K16 to K20 is critical for silencing. In addition, a H4K16Q mutant (which is an acetyl mimic) results in a dramatic loss of silencing (Carmen et al., 2002; Hyland et al., 2005; Lin et al., 2008; Millar et al., 2004; Shahbazian and Grunstein, 2007; Yu et al., 2011) and Sir3 binding is dependent upon the deacetylation of this residue (Ehrentraut et al., 2011; Johnson et al., 1992; Johnson et al., 1990; Onishi et al., 2007; Wang et al., 2013). These data show that the absence of acetyl groups on K16 is crucial for silencing. However, it is currently unknown whether specific nucleosomes have to be unacetylated for silencing or whether a majority of nucleosomes across the entire domain-have to be unacetylated for silencing.

Once established, the silent state is stably maintained for several generations (Gottschling et al., 1990; Pillus and Rine, 1989; Sussel et al., 1993). Occasional disruptions in silencing do occur but are rare: One in a thousand cells stochastically lose silencing at *HML* while ~ seven in ten thousand cells stochastically lose silencing at *HMR*. It is presumed however, that the active state at these loci is short-lived before the silenced state is restored (Dodson and Rine, 2015).

Despite the high fidelity of the inheritance of the silent state, studies suggest that the individual components of silenced chromatin are not stably bound but in constant flux. While the exchange of the core histones in chromatin is quite slow, except at specific regulatory elements (Dion et al., 2007; Misteli et al., 2000), the covalent modifications of the histones have half-lives of only a few minutes (Waterborg, 2001, 2002). While the presence of the Sir3 repressor is essential for silencing (Cheng et al., 1998; Miller and Nasmyth, 1984), analysis of heterochromatin and heterochromatic proteins indicates that repressor protein binding is also dynamic and is influenced by the acetylation and methylation state of the underlying chromatin (Cheng and Gartenberg, 2000; Cheutin et al., 2003; Festenstein et al., 2003). Thus, the overall picture is of a phenotypically stable silenced chromatin state being mediated by constituents that are in constant flux.

Adding further to the complexity of this molecular turmoil is an additional challenge that the cell must overcome to maintain silencing with high fidelity: DNA replication results in a near complete disruption of the chromatin state. Nucleosomes are unable to bind single-stranded DNA (Almouzni et al., 1990) and nucleosomal histones are evicted upstream of the replicating fork (Sogo et al., 1986) and redeposited downstream (Gasser et al., 1996). During DNA replication, nucleosome positions and DNaseI hypersensitive sites (which are sites for binding of transcription factors) are disrupted (Lucchini et al., 2001; Solomon and Varshavsky, 1987; Vasseur et al., 2016) and following replication, the maturation of chromatin leads to the resetting of the original chromatin state (Annunziato and Seale, 1983; Bar-Ziv et al., 2016; Vasseur et al., 2016). The vast majority of the H3/H4 parental tetramers are transferred intact but randomly onto one of the two daughter strands while the parental H2A/H2B dimers segregate randomly to the daughter strands (Annunziato, 2015; MacAlpine and Almouzni, 2013; Mello and Almouzni, 2001). Besides the replication mediated disruption of chromatin structure, the duplication of the DNA also results in the dilution of the parental histone complement by half. The twofold reduction in nucleosome number is restored by newly synthesized histones. Newly synthesized histones are decorated such that histone H4 is acetylated on K5 and K12 and histone H3 is acetylated on K9 and K56 (Benson et al., 2006; Ling et al., 1996; Masumoto et al., 2005; Sobel et al., 1995). The maturation of chromatin following replication involves the removal of these deposition specific modifications of the histones, and the restoration of the modifications found in the mother cell (Bar-Ziv et al., 2016).

The chromatin state that is disrupted during replication, creates a temporal window in the G2 phase of the cell cycle where silenced chromatin is more accessible to enzymatic probes (Aparicio and Gottschling, 1994; Cheutin et al., 2003; Lau et al., 2002) and thus more prone to disruption. Counteracting this disruption are the silencer elements. Elimination of the silencers result in the inability of the silent state to reform following its disruption in S-phase (Cheng and Gartenberg, 2000). Furthermore, efficient inheritability of silencing requires the silencer bound proteins Rap1 and Sir1 (Pillus and Rine, 1989; Sussel et al., 1993).

Besides the silencers, models have invoked a role for histone modification marks in the heritability of the silent state. *In silico* models (Mukhopadhyay and Sengupta, 2013; Sneppen and Dodd, 2012, 2015) suggest that stable inheritance of silencing involves parental modified nucleosomes helping in the templating and modification of nucleosomes containing newly synthesized histones. These models suggest that the efficient inheritance of silenced chromatin likely involves Sir protein binding to unacetylated parental nucleosomes followed by the deacetylation of spatially adjacent newly synthesized histones. The data have also led to a buffer model for the inheritance of the silent state (Huang et al., 2013) which suggests that the silent locus can tolerate significant fluctuations in acetylation levels of the histones during replication. The occasionally acetylated nucleosome at the silent locus does not lead to a loss of silencing but silencing is lost when a particular threshold of acetylation is reached. The level of tolerance in the system is unknown and experiments measuring this are currently lacking. To understand the quantitative relationships between H4K16 acetylation levels and the inheritance of silencing, we developed an assay to quantitatively alter H4K16 acetylation levels and measure the effects of these changes on silencing.

## Materials and Methods

### Protein Blots

Protein lysates were prepared and resolved on a 15% SDS-polyacrylamide gel as described previously (Ghidelli et al., 2001), except that glass beads were used to break open the cells.

### RT-qPCR

Total RNA was isolated from yeast cells as described (Schmitt et al., 1990). cDNA was prepared using the reverse transcription-PCR kit.

### Fluorescence activated cell sorting analysis

Cells were washed in 50mM Tris-HCL, pH7.5 and fixed in 70% ethanol for 1h at room temperature. Cells were then washed in 50mM Tris-HCL, pH7.5 and treated with 1mg/ml RNaseA at 37°C for 1h followed by ProteinaseK treatment (60ug/ml) at 55°C for 1h. Cells were washed and resuspended in phosphate-buffered saline, filtered through a Nitex membrane and stained with Sytox Green stain. Flow cytometry was performed at the UCSC cytometry facility.

#### Fluorescence Microscopy

Cells were grown exponentially in yeast peptone (YP) medium with 2% raffinose at 30°C to an OD_600_ of around 1. The culture was back-diluted to an OD_600_ of 0.125/mL in YP medium with 5 μM alpha-factor and 2% raffinose and incubated on a shaker at 30°C. After 3 hours, the cells were pelleted and transferred into yeast minimal (YM) medium with 5 μM alpha-factor, 2% galactose with appropriate amino acid supplements and incubated on a shaker at 30°C for 4 hours. Cells were pelleted, washed with medium lacking alpha factor, and transferred into YM medium with 2% dextrose and amino acid supplements. Cells were grown on a shaker at 30°C and aliquots removed at appropriate times. After 7 h, the culture was diluted with fresh medium and allowed to grow for another 10 h at 30°C until the final time point.

For each time point, 1 mL of sample was removed and the cells were pelleted and resuspended in 20 uL YM 2% dextrose medium. 3 uL of the suspension was applied to a 1.5% agarose YMD pad on top of a microscope slide and cover-slipped. Images were acquired on a DeltaVision Personal DV system (Applied Precision), using a 40x 1.35 NA oil-immersion objective (Olympus), with a CoolSnap charge-coupled camera (Roper Scientific). 4 μm image stacks were collected, with each Z-image being 0.2 μm apart, 2 μm above and below the plane of focus. Image stacks were taken for each time point and greater than 100 cells were captured across the fields-of-view.

Image analysis was performed using the FIJI distribution of ImageJ software. To measure fluorescence intensity per cell, a two-dimensional maximum-intensity projection was generated for each collected z-stack. A transmitted light image, taken at the center of each z-stack, was overlaid on top of the projection. The transmitted light image served as a guide to establish cell boundaries for maximum-intensity projections, such that maximum fluorescence intensity data could be collected per cell using the software’s measuring tool. Data for approximately 100 cells per time point were collected, compiled into a spreadsheet, and graphed using R software with ggplot2 package.

#### Chromatin Immunoprecipitation

Cells were grown in YPD media to an OD_600nm_ of 2.0 and then fixed with 1% formaldehyde for 10 min. Cells were collected, resuspended in buffer and sonicated using the Bioruptor (Diagenode, Belgium) followed by a cup-horn (Branson, USA) sonicator to an average size of 300bp.

Immunoprecipitation reactions were performed with commercial antibodies to histone H3, Ac-K16 H4 (Millipore, USA), Ac-K56-H3 (Millipore, USA) or with polyclonal anti-Sir3 antibodies and immune complexes were collected with Protein G/A beads (Calbiochem, EMD Biosciences). Immunoprecipitated and input DNA were purified using Chelex 100 (Bio-Rad) (Nelson et al., 2006) and the amount of DNA was quantified using the Picogreeen dsDNA quantitation kit (Invitrogen, USA) and the PerkinElmer Viktor^3^ Fluorescence Reader, prior to qPCR.

Equal amounts of IP DNA and input DNA were used for the qPCR reactions. Quantitative PCR reactions were carried out in a Rotor Gene 6000 with SYBR Green (Platinum SYBR Green qPCR SuperMix UDG, Invitrogen) and a three-step PCR program.

The fold difference between immunoprecipitated DNA (IP) and Input DNA for each qPCR amplified region were calculated as described (Litt et al., 2001), using the formula IP/Input=(2^InputCt − IPCt^). Each experiment involved at least two independent crosslinked samples with each sample immunoprecipitated twice with the same antibody.

## Results

### Histone acetylation is reduced over the silenced domain

We first characterized the chromatin state of the silenced locus in G1-arrested cells to determine the levels of various proteins and modifications at the silenced locus (Figure 1). Using ChIP qPCR, we mapped the relative abundance of core histone H3, Sir3, histone H4K16 acetylation and H3K56 acetylation at the silent *HMR* locus. Mapping histone H3 showed that a site between the two silencers located within the silenced domain had a normal complement of histones as did a site in the euchromatic *GIT1* gene (Figure 1). The silencers as well as the tDNA barrier were moderately “nucleosome-free” as expected though the weaker than expected depletion of nucleosomes is probably due to the average size of the immunoprecipitated DNA (~300 bp) (Cole et al., 2012a; Cole et al., 2012b; Dhillon et al., 2009; Dion et al., 2007; Oki and Kamakaka, 2005).

**Figure 1:**
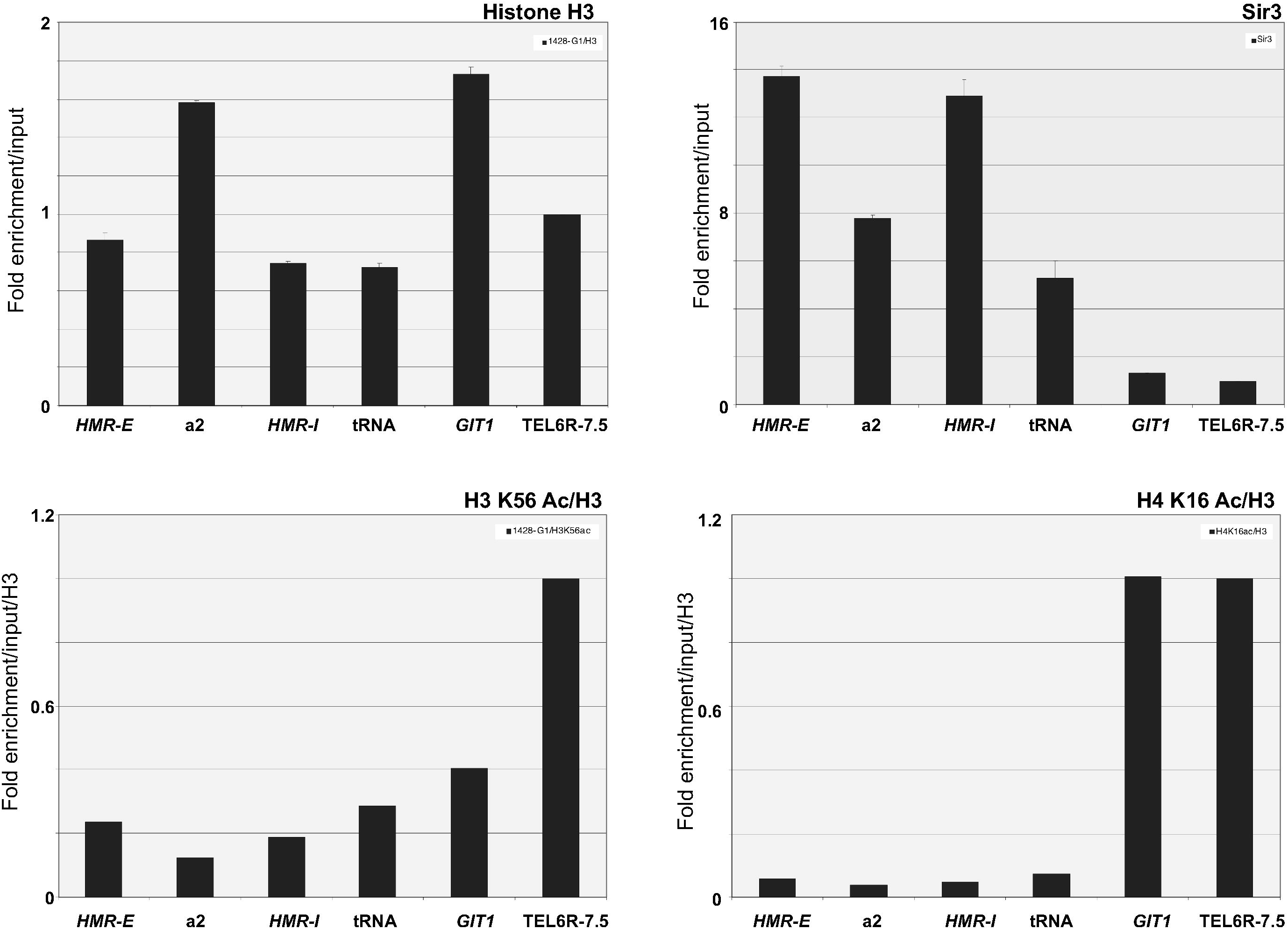
ChIP qPCR of various proteins in G1 arrested cells. Histone H3, Sir3, H4K16 acetylation and H3K56 acetylation levels was measured across the *HMR* domain. Data is presented as the mean enrichment of IP/Input (as described in the materials and methods) for at least four IPs from two independent cross-links. Error bars are standard deviation from the mean. The data for H3K56 acetylation and H4K16 acetylation are presented as enrichment normalized to histone H3 enrichment in order to take into account variable levels of nucleosome occupancy.

Conversely, Sir3 was maximally present at the two silencers while its binding was reduced at the tDNA boundary of the silent domain and at a site within the silent domain which was consistent with previous observations (Thurtle and Rine, 2014; Valenzuela et al., 2008).

We next quantified the distribution of histone acetylation on H3K56 and H4K16 (Figure 1). Since the silencers and the *tDNA* barrier are depleted of histones we normalized the distribution data for these modifications to histone occupancy, thereby showing whether or not there is enrichment or depletion of these modifications on a per-“nucleosome” basis. On a per nucleosome basis, H3K56 acetylation levels showed significant reduction across the entire silent domain and there were even lower amounts of H4K16 acetylation at these sites. The data showed that less than 10% of the histones at the silent locus were acetylated on H4K16 in cells arrested in the G1 phase of the cell cycle.

### Design of the Cut and Flip system

We next investigated the quantitative relationship between histone H4K16 acetylation and gene silencing. Previous work on histones have used one of two different approaches. In one approach, the wild type and mutant histone genes (with their own regulatory elements) are present on plasmids, and the mutant is compared to the wild type strain via plasmid shuffle (Han et al., 1988; Kayne et al., 1988). This system is neither inducible nor tunable and so one is unable to observe the switch or study transition states. In addition, the histone genes are present on plasmids, which fluctuate in copy number from cell to cell. In the second approach, the histone genes are under the control of heterologous enhancer/promoter which can be induced by galactose (Dion et al., 2007). With this approach the histone gene is expressed at high levels throughout the cell cycle in place of its normally restricted expression in the G1/S phase (Eriksson et al., 2012) and this is known to trigger cell cycle checkpoints (Gunjan and Verreault, 2003) and lead to dominant effects (Meeks-Wagner and Hartwell, 1986).

We therefore developed a system to overcome these issues. In *S. cerevisiae* there are two histone loci for the histone H4-*HHF1* and *HHF2*. We constructed a strain lacking the *HHF1* locus and where the wild type histone *HHF2* locus was modified to accommodate two copies of the H4 coding sequence (Figure2A). R recombinase recognition sites flanked the coding region of the wild type H4 gene that had an HA tag at its N-terminus. Immediately downstream of the wild type allele, we inserted a copy of an acetylation mimic mutant of the histone H4 gene fused to an N-terminal Myc tag. This mutant allele lacked the *HHF2* enhancer/promoter element and therefore was not transcribed. The strain also contained the R-recombinase under the control of the *GAL1* enhancer/promoter. The R-recombinase mediated flipping is a rapid and efficient method of creating a desired deletion (Li et al., 2001).

**Figure 2:**
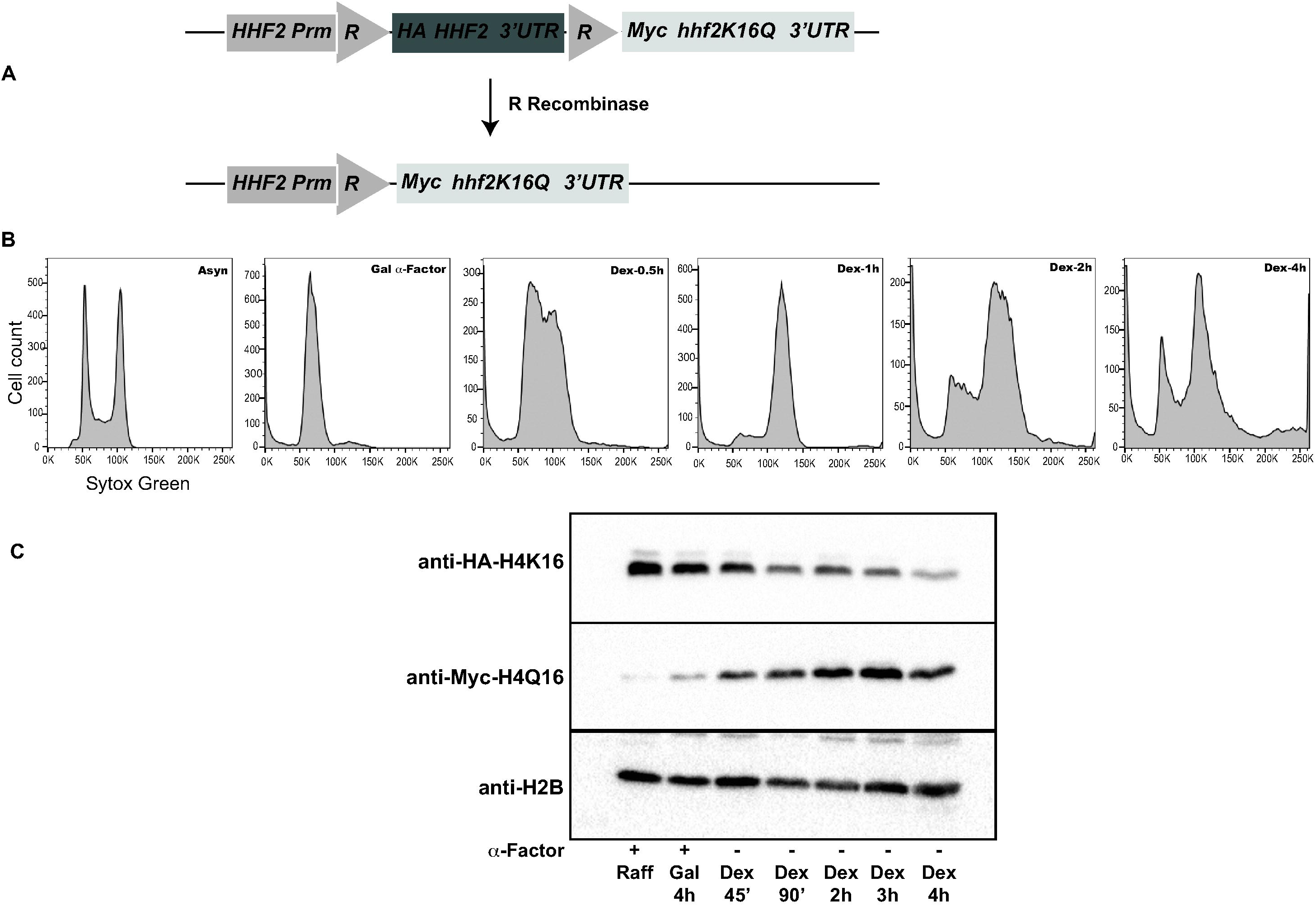
A: Schematic of the Histone H4 cut and flip cassette B: G1 arrest and release fluorescence cytometry profiles of the Cut and Flip strain. Ethanol fixed cells were stained with Sytox Green and analyzed by flow cytometry. Panel 1: Fluorescence cytometry profile of asynchronously growing cells in raffinose containing medium. Panel 2: Fluorescence cytometry profile of cells arrested with alpha factor in galactose containing medium. Panels 3 to 6: Fluorescence cytometry profile of cells at the indicated times after release from alpha factor arrest into glucose containing media. C: Protein immunoblot analysis of cells arrested with alpha factor and released after switching of histone H4 alleles. Yeast cells were grown overnight in raffinose containing rich medium, arrested with alpha factor for 3h and then transferred to galactose containing medium with alpha factor for 4h. Cells were released into YPD and aliquots of equivalent numbers of cells were removed at the specified times. Protein extracts were separated on a 15% SDS-polyacrylamide gel, transferred to membranes and probed with specific antibodies.

The experiment involved growth of yeast cells expressing the wild-type H4 gene from its own UAS enhancer/promoter. Cells were arrested in G1 and the R-recombinase was induced by switching the carbon source to galactose. The recombinase induced recombination between the two R recognition sites flanking the wild type H4 gene which resulted in the flipping out (deleting) of the wild type H4 copy thereby bringing the mutant H4 gene in register with its native UAS enhancer/ promoter. Since the mutant H4 gene is brought under the control of its native UAS enhancer, the mutant protein is expressed only during the G1/S phase of the cell cycle and not over-produced and since the modified histone cassette is present at its native locus on chromosome 14, it does not suffer from changes in copy number.

### Characterization of the Histone H4 Cut and Flip

*MATa* cells (*HML::URA3*p-GFP *GAL1*p-RecR::*LEU2 hhf1-hht1*Δ::KanMx *bar1*Δ::NatMx *HHF2*p-R-HA-*HHF2*-R-Myc-*hhf2K16Q*) were grown overnight in raffinose containing rich medium and arrested in the G1 phase of the cell cycle for 3 hours with alpha factor. We monitored arrest by microscopy (data not shown) as well as by flow cytometry (Figure 2B). Once cells had arrested in the G1 phase of the cell cycle, we shifted the cells to galactose-containing media to induce the R-recombinase. We ascertained that four hours of incubation in galactose were sufficient for maximal R-recombinase mediated switching of the *HHF2* alleles (data not shown). Cells were then released from the G1 arrest into dextrose containing media and aliquots of the cells were removed for further analysis.

Cytometry of the yeast cells (Dhillon et al., 2006) showed that all cells were arrested uniformly in G1. The analysis of these cells following their release from G1 arrest helped us identify the time for each S-phase and showed that the first S-phase occurred around 30 minutes (Figure 2B). The data also showed that most cells progressed through the second S-phase between 2 and 3 hours after their release, albeit with reduced cell-cycle synchrony.

We next monitored the switch of the wild type to mutant *HHF2* alleles by protein blots using antibodies against the HA and Myc epitopes (Figure 2C). Protein extracts were prepared from approximately equal number of cells at each time point and the proteins were resolved on a 15% SDS-Polyacrylamide gel. The proteins after transfer to nitrocellulose membranes were probed with antibodies against HA, Myc or histone H2B. In G1 arrested cells, the predominant histone H4 protein was HA tagged wild type protein. Following release, the levels of histone H4 containing the HA epitope reduced with a concomitant increase in the levels of mutant histone H4-Myc protein. We also monitored the levels of histone H2B as a control and as expected this protein remained relatively unchanged. The protein blots thus demonstrated that the switch cassette functioned as designed.

We then wished to determine if the switched histone H4K16Q mutant proteins were being incorporated into chromatin. Cells arrested in galactose as well as cells collected 2 and 4 hours after release from the G1 phase of the cell cycle were crosslinked with formaldehyde and the crosslinked chromatin was immunoprecipitated using anti-HA and anti-Myc antibodies (Figure 3). Each experiment was performed with a minimum of two independently crosslinked samples and each sample was immunoprecipitated at least twice with the same antibody. The binding of the tagged histones at three different silent loci- *HML*, *HMR* and telomere 6R, was monitored by qPCR. The data showed that the levels of wild type histone H4-HA bound to these loci decreased following release from alpha-factor arrest (Figure 3 top panels), and the levels of mutant histone H4-Myc increased upon release (Figure 3 bottom panels).

**Figure 3:**
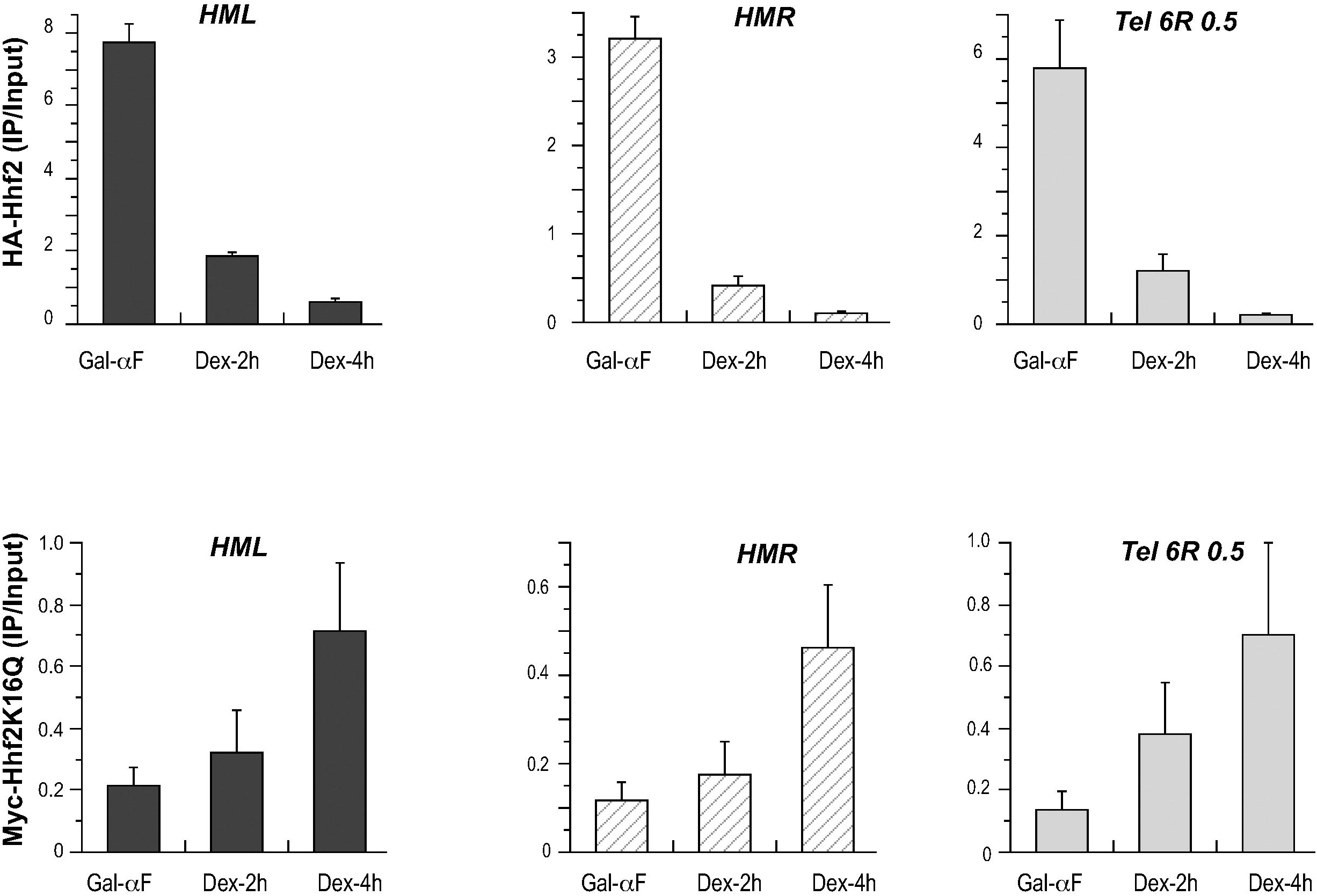
ChIP qPCR of unswitched and switched histone H4 at the silenced loci. The presence of wildtype HA-H4 and mutant Myc-H4 K16Q protein was monitored by ChIP in unswitched and 2h and 4h after switching of the histone H4 allele. The Y-axis represents the ratio of IP/Input DNA for each sample as described in the materials and methods. The levels of the tagged proteins were mapped at three different loci-*HML, HMR* and *TEL6R*.

Having shown that following the switch, mutant histone protein does become incorporated into silenced chromatin, we next investigated the effects of the switch in histones on silenced chromatin using qChIP with antibodies against Sir3 (Figure 4A). In G1 arrested cells, Sir3 was bound to all three silenced loci- *HML*, *HMR* and *TEL6R*. Upon release from the G1 arrest, Sir3 levels reduced within 2h and there was very little Sir3 bound to these loci after 4h showing that incorporation of the mutant histone lead to a loss of Sir3 binding and presumably the activation of the genes at these loci.

**Figure 4:**
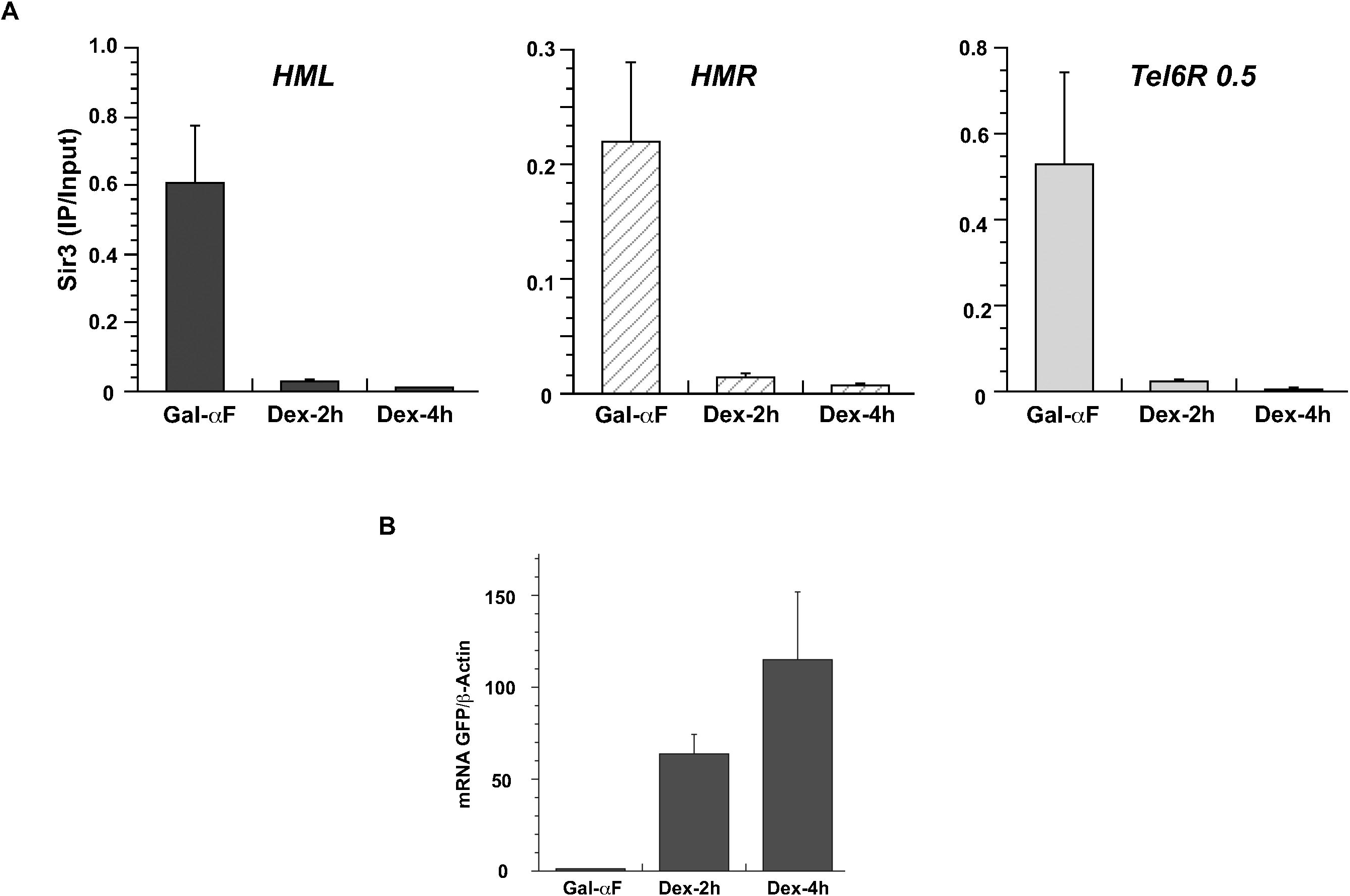
ChIP qPCR measurement of Sir3 binding at silenced loci following switch of WT H4 to H4K16Q mutant A: Sir3 binding at *HML*, *HMR* and *TEL6R* was monitored using ChIP-qPCR in cells arrested with alpha factor and at 2h and 4h after switching the histone H4 allele and alpha factor release. Data is presented as the mean enrichment of IP/Input. B: Measurement of mRNA expression of the GFP reporter at *HML* before and after switch of the histone H4 alleles. Alpha factor arrested cells and cells released into rich medium were collected at 2h intervals and total RNA was extracted from these cells. GFP mRNA was quantitated by RT-qPCR and plotted as a function of time, normalized to *ACT1*.

As a second measure of silencing loss, we measured mRNA levels of a GFP reporter present at *HML* using RT-qPCR (Figure 4B). We isolated mRNA from G1 arrested cells as well as from cells at 2- and 4-hours post-release and measured levels of GFP mRNA along with actin mRNA. In G1-arrested cells there was very little GFP mRNA compared to actin mRNA consistent with the locus being silenced. However, upon release from the arrest, we observed a large increase in GFP expression at the 2h time point which further increased at the 4h time point.

### Fluorescence measurements of gene silencing

Molecular approaches often mask nuance and heterogeneity in data. While one can use mating ability to monitor silencing of the native genes at *HML* and *HMR*, this assesses the silent state only in the G1 phase of the cell cycle and is challenging to monitor in single cells. A fluorescent protein reporter at these loci would circumvent these limitations. We analyzed expression of GFP reporters inserted at *HML*, *HMR* and a telomere using fluorescence microscopy along with the cut and flip cassette. The GFP reporter we employed was a previously characterized, rapidly folding protein (folding/maturation time of ~20min) with a high turnover rate (half-life of ~35min, due to the presence of a *CLN2* PEST sequence) that localized to the nucleus (due to the presence of a nuclear localization signal) (Osborne et al., 2009; Osborne et al., 2011; Xu et al., 2006). We integrated the GFP reporter under the control of either the *URA3* UAS enhancer/promoter or the alpha2 UAS enhancer/promoter at either the *HML* or *HMR* loci or *TEL7L*.

We first built a set of strains expressing either the wild type H4 or H4K16Q mutant protein alone. These strains also contained *HML* and *HMR* loci expressing a GFP reporter under the control of the *URA3* and alpha2 UAS enhancer and core promoter. We measured the GFP signal in at least 100 cells in these strains using a fluorescent microscope (Figure 5A). In cells expressing only the wild type histone H4 protein, we did not observe any GFP fluorescent signal from *HMR::URA3p-GFP, HMR::alpha2p-GFP*, *HML::URA3p-GFP* or *HML::alpha2p-GFP*. In cells expressing only the mutant H4K16Q protein, GFP fluorescence signal was robust and easily detected as predicted for this mutation (Johnson et al., 1990; Lin et al., 2008; Yu et al., 2011). The absolute levels of detected fluorescence in the *H4*K16Q mutant varied both, with the silent locus and the UAS enhancer/promoter. At *HMR*, we consistently saw higher GFP signal when it was under the control of the *URA3* UAS enhancer/promoter compared to the alpha2 UAS enhancer/promoter and we saw a similar expression pattern at *HML*. Interestingly, comparing *HMR* to *HML,* we observed greater derepression of the reporter at *HML* than *HMR,* as well as greater variation in expression of the reporter at *HML* compared to *HMR*. These data suggest that both UAS enhancer/promoter and silencer strength together influence expression levels of the genes at these silenced loci.

**Figure 5:**
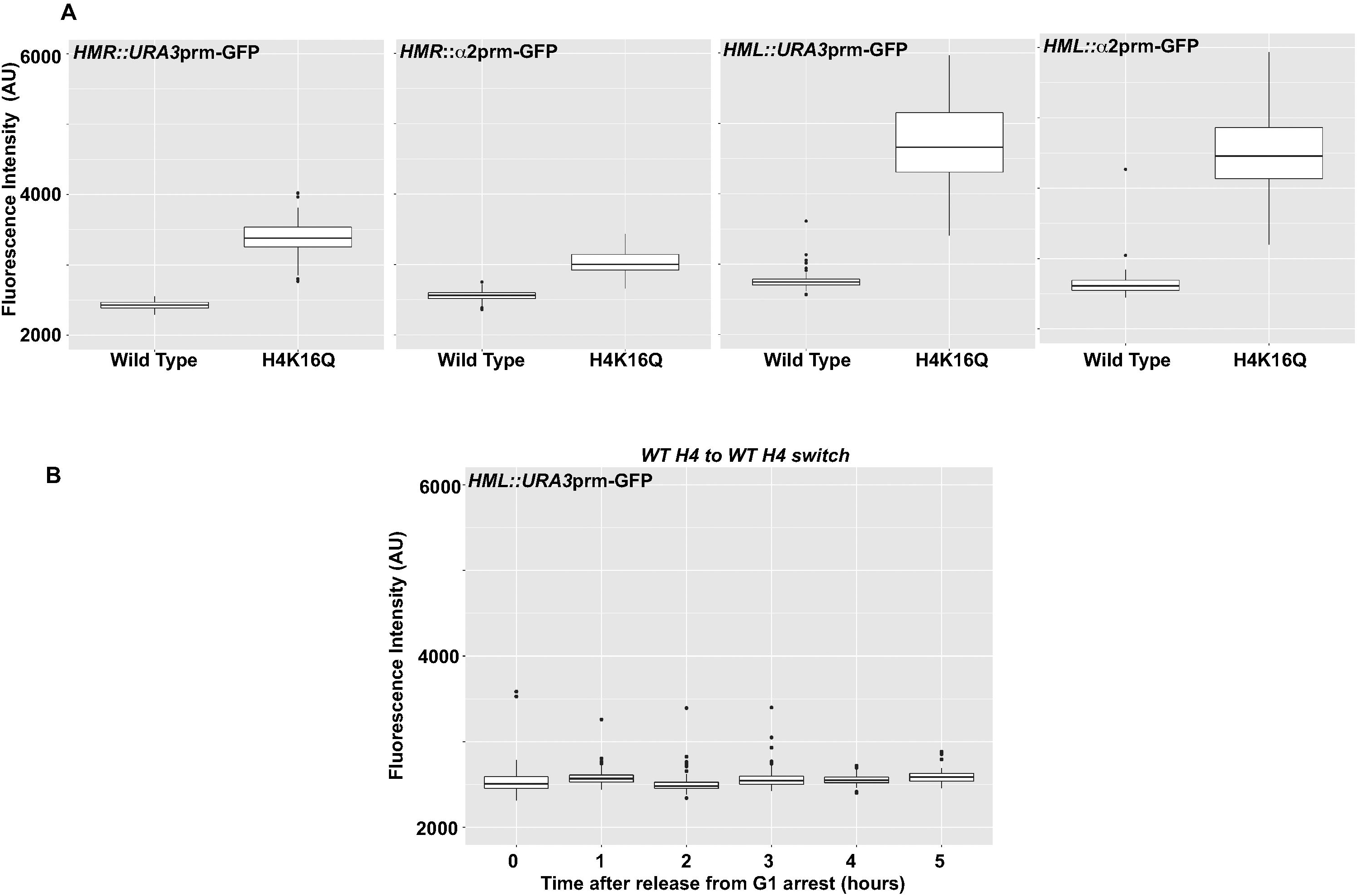
GFP expression in strains with wild type or mutant histone H4. A: Boxplots of GFP expression from silenced loci in strains expressing WT and mutant histone H4 K16Q alleles Cells were grown in rich medium, imaged using a fluorescence microscope and the amount of fluorescence in each cell was quantitated and plotted as a box plot. For each sample, GFP fluorescence was measured in greater than 100 cells. B: Fluorescence measurements of GFP at *HML* following switching the histone H4 cassettes A wild type histone HA-H4 cassette was switched to a wild type Myc-H4 cassette in G1 arrested cells and silencing at *HML::GFP* was monitored in the cells after their release from the cell cycle arrest.

We also wished to confirm that the act of switching the histones did not perturb the silent state. We generated a cut and flip *HHF2* strain where the wild type H4 could be switched to another wild type H4 (*HHF2*p-R-HA-*HHF2*-R-Myc-*HHF2)*. Cells were arrested in G1, the cassette was switched and then cells were released into the cell cycle. GFP expression at *HML::URA3p-GFP* was then measured over time (Figure 5B). We did not observe any changes in GFP fluorescence upon switching of the histones; therefore, the histone switch in and of itself did not affect silencing.

To determine the quantitative relationship between H4K16Q levels at the silent loci and gene silencing, we employed strains where the wild-type H4 could be switched to a mutant H4K16Q. We arrested these cells in G1, switched the histone alleles using R-recombinase and then released these cells from the G1 arrest and monitored expression of GFP by fluorescence microscopy. At *HML,* when GFP was under control of the *URA3* UAS enhancer/promoter, measurable fluorescent signal was observed 2h after release from G1 arrest and reached maximal levels around 5h. These data suggest that silencing was beginning to be lost during or soon after the second S-phase (Figure 6A).

**Figure 6:**
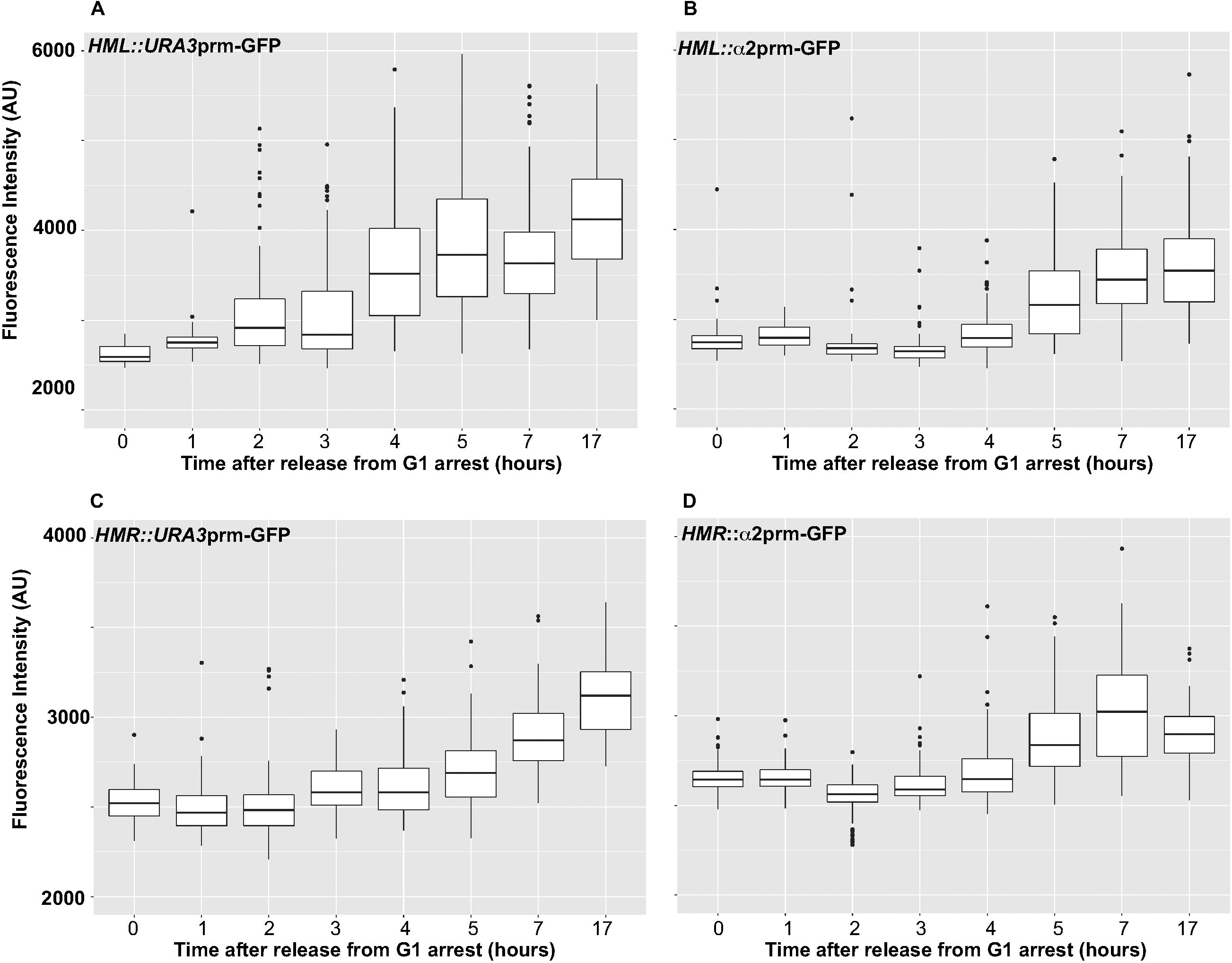
Boxplots of GFP expression at *HML* and *HMR* as a function of time after switching the histone H4 cassettes GFP fluorescence was measured as a function of time in strains with modified *HML* and *HMR* loci containing GFP under the control of either the *URA3* UAS enhancer/promoter or the alpha2 UAS enhancer/promoter. Cells were arrested in G1, the histone H4 cassette was switched from wild type H4 to mutant H4K16Q and cells were the released from the arrest. GFP fluorescence was measured as cells progressed through the cell cycle.

When we measured GFP expression under the control of the alpha2 UAS enhancer/promoter at *HML*, measurable fluorescence was first observed around the 4h time point with maximal expression occurring around the 7h time point indicating that silencing was beginning to be lost in or after the third S-phase (Figure 6B).

We saw similar dynamics for the *HMR* locus. When the GFP reporter was under the control of the *URA3* UAS enhancer/promoter, we saw measurable GFP signal approximately 3h after the release while for the alpha2 UAS enhancer/promoter, GFP signal was first observed 4h after the release (Figure 6C and 6D).

We also analyzed silencing at telomere 7L. The GFP reporter under the control of the *URA3* UAS enhancer/promoter was inserted adjacent to *TEL7L*. Cells were arrested in G1, the histone allele was switched and GFP expression was measured after release. A measurable fluorescent signal was observed within 1h after release suggesting that ~50% replacement of wild type H4 with H4K16Q was sufficient for weakening the silent state at this locus (Figure 7A).

**Figure 7:**
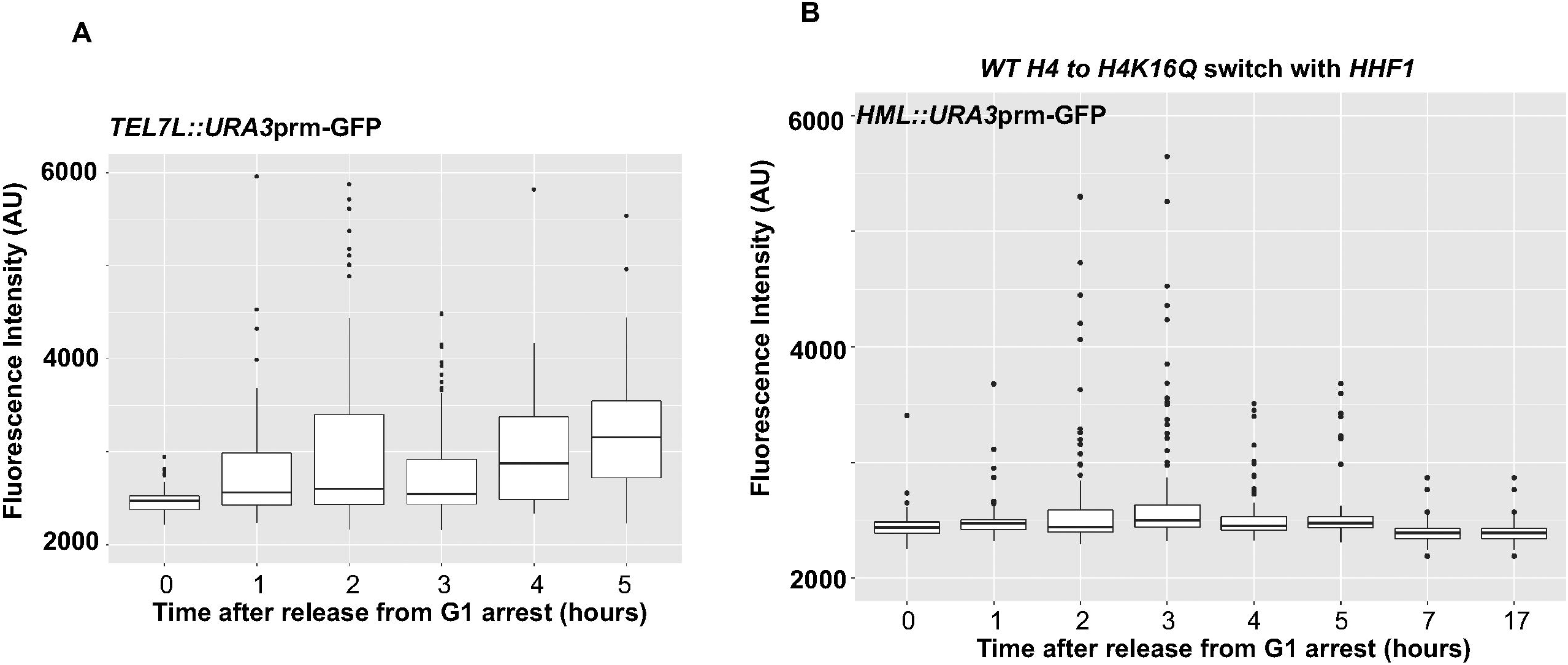
Boxplots of GFP expression at the Telomere and *HML* following switching the histone cassette A: GFP fluorescence measured as a function of time in strains with *TEL7L::URA3p-GFP.* B: GFP fluorescence was measured as a function of time in strains with *HML::URA3p-GFP* but also containing the wild type copy of the *HHT1-HHF1* locus.

It is possible that for *HML* and *HMR*, silencing in some cells begins to be lost at early time points but the increases in expression went undetected due to the limitations in the sensitivity of our fluorescent measurement set up. We nevertheless observed quantifiable loss-of-silencing at *TEL7L* at these early time points, showing that the telomeres are more susceptible to changes in histone acetylation than the cryptic mating type loci and the inability to detect GFP signal from *HML* and *HMR* at early time points is not due to the time required for the maturation of the GFP fluorescent signal.

In this study we quantified silencing by measuring levels of GFP fluorescent signal in individual live yeast cells. The actual time when silencing is lost and transcription initiates from the silent locus will be different from the time when GFP fluorescent signal is detected by microscopy. The GFP mRNA is ~1000 bases long and with a yeast transcription elongation rate of 25 bases/second (Pelechano et al., 2010) would be transcribed within ~40 seconds. The yeast translation rate is 2.63 amino acids/second (Riba et al., 2019) and so GFP would be translated in ~2 minutes. The maturation time of the GFP protein used in this study is ~20 minutes (Osborne et al., 2009; Osborne et al., 2011; Xu et al., 2006) and thus detection of the GFP fluorescent signal would be delayed ~23 minutes from the actual time of loss of silencing. Since we used one-hour time points for our fluorescence measurements, we do not believe that this offset prevents us from correlating our observations to cell cycle events.

Our results showed that at *HML* and *HMR,* silencing was not lost after the first S-phase but weakened during or after the second S-phase, when the wild type H4 levels should have dropped to at least 25%. To confirm this result we built a cut and flip *HHT2-HHF2* strain that contained the wild type *HHT1-HHF1* alleles, thereby halving the fold-reduction of the wild type H4 with each DNA replication event. In this strain, the percent of chromatin-bound H4K16Q would approximately be 25% after the first S-phase, increase to 37.5% after the second S-phase and approach 50% after successive S-phases. We arrested this strain in G1, switched the *HHF2* allele from wild type to H4K16Q, and monitored expression of the *URA3* UAS enhancer/promoter driven GFP reporter at *HML* (Figure 7B). In this strain, we did not observe expression of GFP after switching the *HHF2* alleles from wild type to mutant suggesting that greater than 50% H4K16Q histones need to be incorporated at *HML* before a quantifiable GFP fluorescent signal can be observed.

## Discussion

The silencer and silencer bound proteins are necessary for efficient inheritance of the silent state (Cheng and Gartenberg, 2000; Pillus and Rine, 1989; Sussel et al., 1993). The key role of the silencers is to maintain a high concentration of Sir proteins in the vicinity of the locus for the state to be re-established after its disruption during replication. It is likely that silencer strength influences the efficiency of inheritance since we consistently observe greater silencing mediated by the *HMR* silencers compared to the *HML* silencers in agreement with previous observations about silencer strengths (Shei and Broach, 1995).

In addition to the silencer, efficient inheritance of the silent state depends upon the nucleosomes remaining unacetylated. There are approximately 20 and 12 nucleosomes present at *HML* and *HMR* respectively (Ravindra et al., 1999; Weiss and Simpson, 1998). While it is possible that the deacetylation of a single key nucleosome is necessary for silencing, our data argue against this. We support a model where the locus requires an aggregate level of acetylated nucleosomes for silencing to be lost. In this scenario, a domain would remain silent so long as the number of unacetylated nucleosomes are above a certain threshold. The silent locus can thus tolerate fluctuations in overall acetylation levels without functional consequence. Our data suggest that for *HML* and *HMR* to lose silencing, around 75% of the nucleosomes must acquire acetyl marks before the locus loses silencing. That acetylation of 50% to 75% nucleosomes causes loss of silencing is also consistent with biochemical data showing that Sir proteins are able to interact with unacetylated histone tails in one nucleosome and bridge neighboring nucleosomes leading to silencing (Behrouzi et al., 2016; Ehrentraut et al., 2011; Ghidelli et al., 2001; Johnson et al., 1992; Onishi et al., 2007; Wang et al., 2013).

The bulk of the yeast nucleus is packaged into euchromatin and consistent with this is the observation that almost every histone H4 molecule is acetylated (Hecht et al., 1995; Kuo et al., 1998; Waterborg, 2001). The exception to this is the silent loci where histone H4 molecules are not acetylated. If one assumes for simplicity’s sake that H4K16 acetylation is required for the spontaneous loss of silencing, then our data can be used to calculate the probability of a stochastically spontaneous acetylation of a nucleosome at the silent locus. At the native silenced loci, silencing is stochastically lost in one out of every 1000 cells at *HML* with a similar value at *HMR* (Dodson and Rine, 2015). Based on our data, for one cell out of 1000 to spontaneously lose silencing, 75% of the nucleosomes would need to acquire H4K16 acetylation in that cell. Therefore, at *HML*, for 15 out of the 20 nucleosomes (75%) to be simultaneously acetylated, a single nucleosome would have a ~1/1.6 (60%) probability of acquiring an acetyl group by chance. These numbers suggest that just a small reduction in the ability of acetylases to acetylate nucleosomes across a contiguous stretch of DNA may be sufficient to generate a transcriptionally silent region in the nucleus. The precise number would likely vary from cell to cell since other factors such as the local concentration of the Sir proteins, transcription activators and modifications of other histone residues (such as H3K56 and H3K79) are also likely to fluctuate and affect the overall process.

### Replication and acetylation

Silencing is a dynamic equilibrium state and the key determinants for resetting the silent domain following replication would be the relative local concentrations of transcription activators (and coactivators) and repressor (and corepressor) proteins at these loci (Aparicio and Gottschling, 1994; Donze et al., 1999; Renauld et al., 1993; Shei and Broach, 1995; Valenzuela et al., 2009). While silencing is mediated by proteins in constant flux, it is nevertheless stable and faithfully propagated through growth and cell division. There are likely many different factors that collectively lead to this high fidelity. The parental histones segregate randomly to the replicated daughter strands and in theory parental histones with active modifications (such as H4K16 acetyl) could ingress into the silenced domain and aid in the switch from silent to active state. However, while parental histones are evicted from the DNA during replication, they are re-deposited in close proximity to their original site, thereby reducing the probability of histones with active modifications being transferred to silenced chromatin (Jackson and Chalkley, 1985; Radman-Livaja et al., 2011). Moreover, active chromatin is replicated early while silenced loci are replicated late (Friedman et al., 1995; Raghuraman et al., 2001) and this temporal separation would further reduce the likelihood that silent loci would become bound by parental histones containing active chromatin marks. It is also highly unlikely that silent loci acquire acetyl marks from newly synthesized histones, since newly synthesized histone H4 is acetylated on K12 and not K16 (Ai and Parthun, 2004; Sobel et al., 1995). In addition, the presence of the silencers increases the local concentration of the Sir proteins compared to the global nuclear distribution of Sas2 acetylase throughout the nucleus, thus reducing the probability of nucleosome acetylation and favoring the deacetylated state at silent loci. Lastly, the three-dimensional clustering of silent loci (Kirkland and Kamakaka, 2013; Maillet et al., 1996) could create a pinball effect, trapping Sir proteins in the vicinity of the silent loci and increasing the effective local concentration of the Sir proteins at these loci. While Sir2 removes acetyl groups from nucleosomes that stochastically acquire the modifications because of the global presence of Sas2, the primary function of the other Sir proteins is likely the prevention of acetylation of the histones following their deposition onto newly replicated DNA.

### Binary versus Analog silencing

Silencing has classically been shown to be an all-or-nothing phenomenon: a locus is either fully silent or active (Gottschling et al., 1990; Pillus and Rine, 1989). An interesting observation from our studies is that during the loss of silencing at early time points we did not observe a digital “binary” response in the levels of GFP protein. When we measured the amount of GFP fluorescence in individual cells, we observed a continuum of values. If one assumes a proportional relationship between mRNA levels and protein levels, and also assumes that the expression of the mating type transcription factors is similar to GFP expression, then the levels of GFP observed in hundreds of individual nuclei do not show a bimodal distribution suggesting that loss of silencing was not an all-or-nothing phenomenon and in a population of cells loss of silencing generates a continuum of mRNA (Dodson and Rine, 2015) and protein levels that, at the level of a phenotype, (ability to mate) translates into a binary choice-mating or non-mating.

The implications of a partially silent state are variable levels of GFP proteins in individual cells at these early time points. One possibility to explain this phenomenon is that transcription is noisy and occurs in bursts. If one assumes that transcription is a probabilistic event then partial silencing could be due to changes in transcription burst frequency or burst size (Otto, 2019; Rodriguez and Larson, 2020; Wang et al., 2018). This, in turn, would be determined by accessibility of the UAS enhancer and core promoter to the transcription machinery. Access could be affected by altering the affinity of the transcription factors for their binding sites via post-translational modifications of these proteins or by the modifications of nucleosomes leading to changes in the positioning of nucleosomes over the binding sites as well as by altering the ability of chromatin remodelers to evict or slide nucleosomes. Thus, the role of the Sir proteins in transcriptional silencing would be to alter the probability of transcription activator and general factor binding to the UAS enhancer and core promoter, as well as alter the ability of chromatin remodelers to slide or evict nucleosomes.

## Acknowledgements

This work was supported in part by a grant from the NIH to RTK (GM078068) and (T32-GM008646) to KW. We would also like to thank J. Rine for providing us with specific yeast strains. We would like to thank N. Bhalla for the use of her fluorescence microscope and the UCSC cytometry facility funded by the CIRM Shared Stem Cell Facility grant to UCSC (CL1-00506) for help with fluorescence cytometry.

